# A novel score for highly accurate and efficient prediction of native protein structures

**DOI:** 10.1101/2020.04.23.056945

**Authors:** Lu-yun Wu, Xia-yu Xia, Xian-ming Pan

## Abstract

Protein structure resolution has lagged far behind sequence determination, as it is often laborious and time-consuming to resolve individual protein structure – more often than not even impossible. For computational prediction, due to the lack of detailed knowledge on the folding driving forces, how to design an energy function is still an open question. Furthermore, an effective criterion to evaluate the performance of the energy function is also lacking. Here we present a novel knowledge-based-energy scoring function, simply considering the interactions of peptide bonds, rather than, as conventionally, the residues or atoms as the most important energy contribution. This energy scoring was evaluated by selecting the X-ray structure from a large number of possibilities. It not only outperforms the best of the previously published statistical potentials, but also has very low computational expense. Besides, we suggest an alternative criterion to evaluate the performance of the energy scoring function, measured by the template modeling score of the selected rank-one. We argue that the comparison should allow for some deviation between the x-ray and predicted structures. Collectively, this accurate and simple energy scoring function, together with the optimized criterion, will significantly advance the computational protein structure prediction.

---

A protein’s structure determines its thermodynamic and biological function, and provides the three-dimensional basis for drug design^1-3^. However, the resolution of protein structures lags far behind their sequence determination, since the resolution often is laborious, time-consuming, and sometimes impossible^4^. To facilitate the resolution process, computational prediction methods have been developed. However, the accuracy of the predicted structures is still being questioned^5-7^.

The total size of the conformational landscapes increased exponentially with the increasing of the protein chain length and very quickly become astronomical^8^. Designing an accurate and simple energy scoring function is a crucial task in protein structure prediction. Several physics- and knowledge-based energy functions have been developed. Physical energy functions such as CHARMM, AMBER, GROMOS, and OPLS^9-12^ are widely used in areas such as protein structure refinement. However, they have limited application in protein structure prediction, due to the computational expense and lack of significant gains^13,14^. As an alternative to physics-based energy functions, knowledge-based energy functions have been widely employed in protein structure prediction^15-18^. However, according to the CASP, no method is able to predict the structures of all kinds of proteins. Here, we assume that the interactions between a protein’s peptide bonds, rather than as conventionally, the residues or atoms are the most important energy contribution. We design a novel energy scoring function, named PBE scoring function. The PBE scoring function not only outperforms the tops of the previously published energy scoring function, but also has a factor of 39 to 350 lower CPU time requirement. Additionally, the parameters of the PBE scoring function were obtained from fitting an in-house dataset, where each protein has a sequence identity less than 20% with that in all test datasets used to avoid over-fit problem. Our results suggest that peptide bond is much more suitable for constructing knowledge-based energy functions than residues or atoms. This energy function should advance the computational protein structure prediction and highlight the importance of the interactions between peptide bonds as the protein folding driving force.

Another challenge for computational structure prediction is to identify when the prediction is correct. Currently, the predicted structure is considered correct only if it exactly matches the experimentally determined x-ray structure. However, we believe this criterion is unjustified: Often, we find that predictions are classified as erroneous when (1) there is uncertainty in the x-ray structure, or (2) the difference between the x-ray and the predicted structures is so small that the two structures are identical for all practical purposes. Thus, we propose a novel criterion that takes these challenges into account. With a structural-similarity threshold TM-score >0.7, the PBE scoring function achieves a success rate of 100% on the CASP5-8 dataset

In the following, we will first present the design of the PBE energy function and the PBE’s prediction performance with the conventional criterion. Then, we present the new criterion, before re-evaluating the PBE’s prediction performance with the new criterion.

## Design of the PBE function

To choose the native structures from a large number of decoys, we design a new protein PBE scoring function. We focus on the peptide bond, since proteins are composed of amino acids or residues joined together by peptide bonds. Note that the peptide bond has a slight double-bond character (40%). Due to resonance, which occurs with amides, the C-N bond length is 10% shorter than that found in usual C-N amine bonds. Furthermore, C_α_, C and O atoms of the *i*th residue, as well as C_α_, N and H atoms of the (*i+1*)th residue, are approximately coplanar. This rigidity of the peptide bond reduces the degrees of freedom of the polypeptide during folding. As a consequence, all peptide bonds in protein structures are found to be almost planar. This planar characteristic of the peptide bond constrains the protein rotation, which finally shapes the overall three-dimensional protein structure. The interaction with one residue (or atom) within a peptide bond would be conducted to all groups within this peptide due to the resonance effect, indicating that the interaction with the group within a peptide bond is nearly equivalent to the interaction with the entire peptide bond. For these reasons, our method focuses on the protein peptide bonds and their directions, in which the proteins’ peptide bonds are considered as the structural entities,

In a protein structure, the *i*th residue is represented by the coordinate vector *D*_*i*_ of its *C*_*α*_-atom, and then its peptide bond direction is defined by the vector *V*_*i*_ (*V*_*i*_ = *D*_*i*+1_ − *D*_*i*_). The interaction between the *i*th and *j*th peptide bonds is represented by the cosine value (cos(*θ*): *θ* is the angle between *V*_*i*_ and *V*_*j*_). To calculate the PBE score of the *i*th peptide bond, we consider 3 adjoining peptide bonds, namely, *i*-1, *I* and *i*+1, and assume that it is affected by both localized and distant interactions The PBE score of a peptide bond is defined as:(eq. 1):

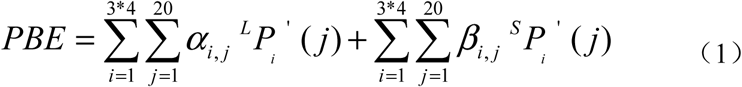

The PBE score of a structure is defined as the average of all peptide bonds of the structure. In equation 1, the first term on the right hand side describes the long-range peptide interactions, while the second term denotes the short-range peptide interactions.

The long-range term represent the interactions between the peptide bonds in the window, and the spatially closest bonds but in sequence outside the window. Three long-range interactions occur for each peptide bond in the window, producing 3*4=12 new PSSM matrix vectors *L* _*Pi*_’, where α_i,j_ denotes the coefficients. The second item represents the short-range interaction between the three peptide bonds in the window. Three short-range interactions also exist, producing 3*4=12 new PSSM matrix vectors *S* _*Pi*_’, where *β*_*i, j*_ denotes the coefficients. In total, 480 coefficients exist in the energy scoring function.

We assume that the interaction between two peptide bonds leads to a change in the values of the PSSM matrix vector *P* of the four residues included in these two peptide bonds.

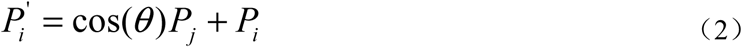

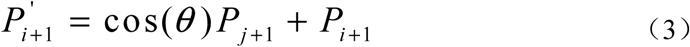

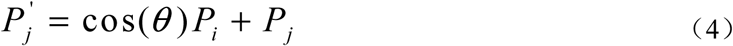

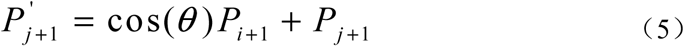

Furthermore, we classify the PBE score function into 8*8*8=512 categories according to the cosine values of the short-range interactions. In each category, the samples of the in-house dataset can be used to set up more than 600000 equations to fit 480 parameters. This large number of training samples ensures the validity of parameters and avoids the over-fitting problem.

All the coefficients are determined by the in-house training data set using a multiple linear regression method to minimize the sum of the square of the deviations between the left and the right side of the equation (1)^19^. During training, for each peptide bond, the PBE score on the left side of equation (1) is set to 0.0 for an x-ray structure and to 1.0 - TM-score for a decoy structure. The prediction is performed with “Jackknife” analysis^19^, exploiting the database. Each of the sample proteins in the database (including the x-ray structure and decoys) is excluded from the calculation of the coefficients. These coefficients are used in equation (1) to calculate the PBE score of the excluded proteins. Under the self-consistent validation, the PBE function score correctly chooses all 5147 x-ray structures as having the lowest free energy, achieving a success rate of 100% and a Z-score of -9.67. When we perform the jackknife test, 5070 protein X-ray structures are correctly identified as having the lowest free energy. This performance equals a success rate of 98.62% and a Z-score of -9.01 (Table 1 in extended data). The success rate decreases only by 1.4% under the jackknife test, suggesting that the score is quite robust.

**Table 1.** Comparison of the *PBE* with other energy function scores.

| Function | Rank $\leq 5$ | Rank = 1 | z-score | Cost of<br>CPU-time/s |
| --- | --- | --- | --- | --- |
| **PBE** | **140** | **122** | **-2.84** | **19** |
| **KBF** | 138 | 97 | -1.92 | 6701 |
| **RW** | 136 | 110 | -1.69 | 915 |
| **RWplus** | 135 | 106 | -1.69 | 2010 |
| **4BOPT** | 131 | 72 | -1.46 | / |
| **dDFIRE** | 128 | 100 | -1.41 | 743 |
| **SKOb** | 125 | 48 | -1.45 | / |
| **fourBody** | 124 | 53 | -1.30 | / |
| **SKOa** | 122 | 49 | -1.27 | / |
| **SKJG** | 120 | 55 | -1.37 | / |
| **4BOPT_GNM** | 119 | 46 | -1.06 | / |
| **MJ3** | 114 | 46 | -1.36 | / |
| **HLPL** | 113 | 50 | -1.28 | / |
| **Qp** | 112 | 57 | -1.32 | / |
| **BFKV** | 112 | 57 | -1.52 | / |
| **MS** | 112 | 42 | -1.34 | / |
| **VD** | 111 | 53 | -1.37 | / |
| **TEs** | 111 | 46 | -1.39 | / |
| **Qm** | 111 | 42 | -1.21 | / |
| **Qa** | 111 | 37 | -1.16 | / |
| **genFour** | 110 | 53 | -1.25 | / |
| **Tel** | 108 | 46 | -1.36 | / |
| **shortRange** | 108 | 39 | -1.05 | / |
| **BT** | 106 | 58 | -1.49 | / |
| **MJ3h** | 106 | 51 | -1.34 | / |
| **TD** | 105 | 57 | -1.30 | / |
| **GKS** | 103 | 38 | -1.18 | / |
| **MJPL** | 100 | 45 | -1.02 | / |
| **MJ2h** | 96 | 45 | -1.05 | / |
| **TS** | 95 | 36 | -1.00 | / |
| **MSBM** | 61 | 17 | -0.11 | / |
| **RO** | 57 | 7 | -0.36 | / |
| **MJ1** | 14 | 0 | 0.76 | / |

/: not calculation

## Performance on conventional criterion

To measure the performance of the PBE function, we try to predict the exact x-ray structures in the widely used CASP5-8 dataset. This dataset contains 2628 possible structures (*i*.*e*. decoys) for 143 proteins. The PBE function assigns the lowest free energy to the exact x-ray structure for 122 of the 143 (85%) proteins, and achieves a Z-score of -2.84. The 85 % and the Z-score are both higher than for all 32 considered, previously published energy functions (table 1). The next highest performance is achieved by the functions RW, RWplus, dDFIRE, and KBF, which identified 110(77%), 106(74%), 100(70%) and 97(68%) x-ray structures respectively. Also the cost of CPU time of PBE is significantly lower than all the previously published functions. *E*.*g*., the cost of CPU time on the CASP5-8 dataset is 19 seconds for the PBE, while 743 seconds for dDFIRE, 915 seconds for RW, 2010 seconds for the RWplus and 6701 seconds for KBF. Thus, the PBE constitutes a significant improvement in both computational expense reduction and accuracy improvement.

## The novel evaluation criterion

Our second contribution, to the improvement of computational protein structure prediction, is the novel prediction evaluation criterion. To develop this criterion, we investigate the reason why the PBE’s predictions were classified as failures for 21 proteins in the CASP8-5 dataset. For this prediction, we employ the criterion where we require the exact match with the x-ray structure. Thus, we investigate the difference between the 21 x-ray and the 21 PBE predicted structures: In detail, we evaluate these structural differences with the template modeling score (TM-score)^20^, as the TM-score measures the global similarity of two protein structures and is length-independent. According to Zhang, *et al*.^21^, two protein structures with a TM-score > 0.5 are likely to have the same topology, while those with a score > 0.8 certainly have the same topology. Interestingly, when the TM-score is greater than 0.9, both Zhang and Hrabe^22^, *et al*. suggest that it is unnecessary to distinguish the predicted structures from their corresponding x-ray structures, since these two structures are highly likely to be identical. Furthermore, when the TM-score between predicted and x-ray structure is above 0.5 to 0.7^21^, the predicted structure is sufficiently accurate to be used in drug design. Within the 21 proteins where the PBE prediction was classified as an erroneous when requiring the exact x-ray structure, there are 6 structures with a TM-score greater than 0.9, 9 structures with a TM-score between 0.8 and 0.9, and 6 structures with a TM-score between 0.7 and 0.8. Thus, there are 6 proteins where the predicted structure is, for all practical purposes, identical with the x-ray structure (TM score > 0.9). For these 6 proteins, we still judge the PBE to have failed since we require the exact match between predicted and x-ray structure. Moreover, *all* the 21 structure predictions would have been sufficient for use in drug design, as their TM scores were above 0.7, but they were still classified as erroneous. Thus, by requiring the exact match, insignificant differences to the x-ray structure can lead to the prediction being classified as a failure. In our opinion, these predictions should not necessarily have been classified as a failure since they were either virtually identical to the x-ray structure (TM score > 0.9) or accurate enough for some task (TM score > 0.7). This motivates us to shift away from requiring the exact match with the x-ray structure as the prediction criterion.

Another reason not to require the exact match between the predicted and x-ray structure is that there can be experimental uncertainty in the x-ray determined structure: *E*.*g*., Hrabe, *et. al*. reported that the root mean square deviation (RMSD) between x-ray structures of the same protein may range from 0-4Å^22^, when the structure is determined by different groups. These 0-4 Å stem from differences in apo and holo forms, physicochemical-condition variations during crystallization, range of the conformational ensemble of the individual protein, etc.. Even, it is possible that the x-ray-determined structure is less similar to the native structure than a predicted structure^23^. Figure 1 shows the relationship between the TM-score and the average RMSD of the high-resolution training set^24^. At a TM-score of approximately 0.7, the average RMSD is approximately 3Å, which decreases with increasing TM-score. At a TM-score of approximately 0.9, the average RMSD value is approximately 2Å, indicating that structures with TM-scores greater than 0.9 have the same topology as that of the X-ray structure. We further investigate this x-ray structure uncertainty and the divergence with the predicted structures, by using MaxCluster ^25^. MaxCluster aligns the 21 predicted with the 21 x-ray structures. When aligned, there is a weak correlation between the temperature factors (B-factor) (Figure 2 presents 3 examples; extended data Figure 1 presents all 21 proteins) and their corresponding *C*_*α*_ RMSD values. Thus, in some local regions where the B-factors are high, the *C*_*α*_ RMSD values are also high. *I*.*e*., the correlation between high B-factor and high RMSD value suggest that the x-ray and predicted structures mainly diverge in the region where the protein is flexible. In the areas where the protein is not flexible, the prediction exactly matches the x-ray structure. Indeed, x-ray structure determination of a flexible protein can be difficult, as the x-ray method generally detects the average conformation of proteins, which may not have the lowest energy^6,26,27^. Thus, our analysis of the 21 proteins and the literature agree, that requiring the exact match with the x-ray structure suffers from experimental uncertainty in the x-ray structure.

**Figure 1.**
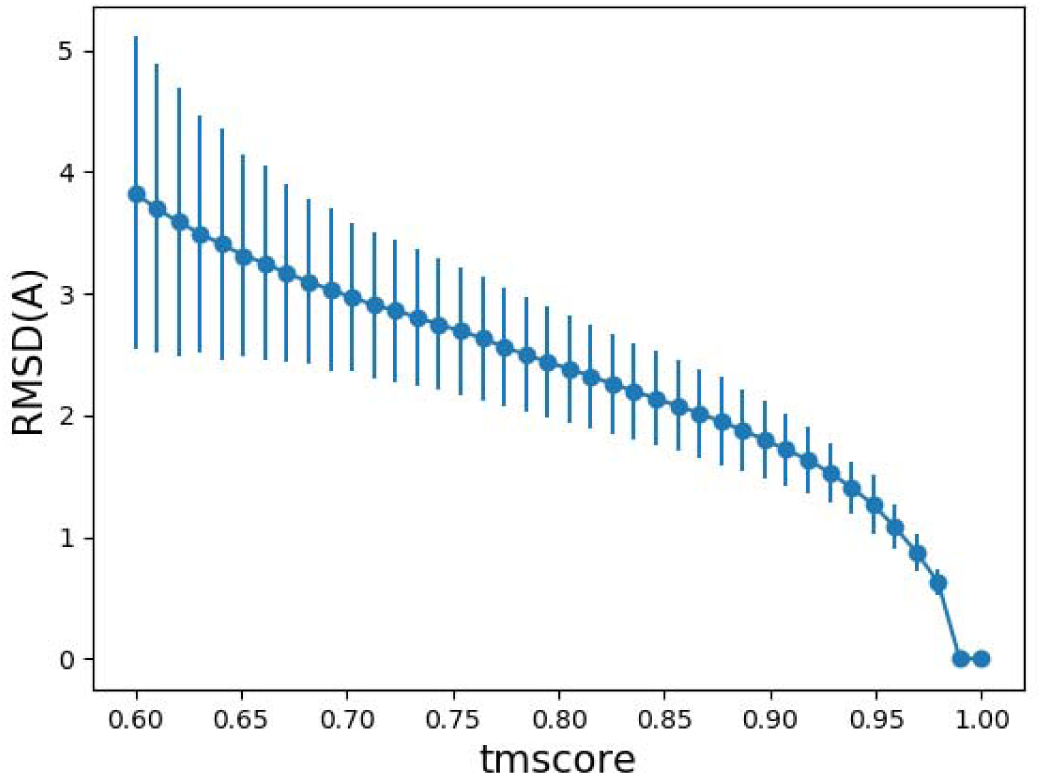
The relationship between the TM-score and the average RMSD of the high-resolution training dataset. The average RMSD and the standard deviation of RMSD decrease sharply as the TM-score increases. When TM-score >0.4. the average RMSD is below 3 Å and the standard deviation of RMSD is below 1 Å.

**Figure 2.**
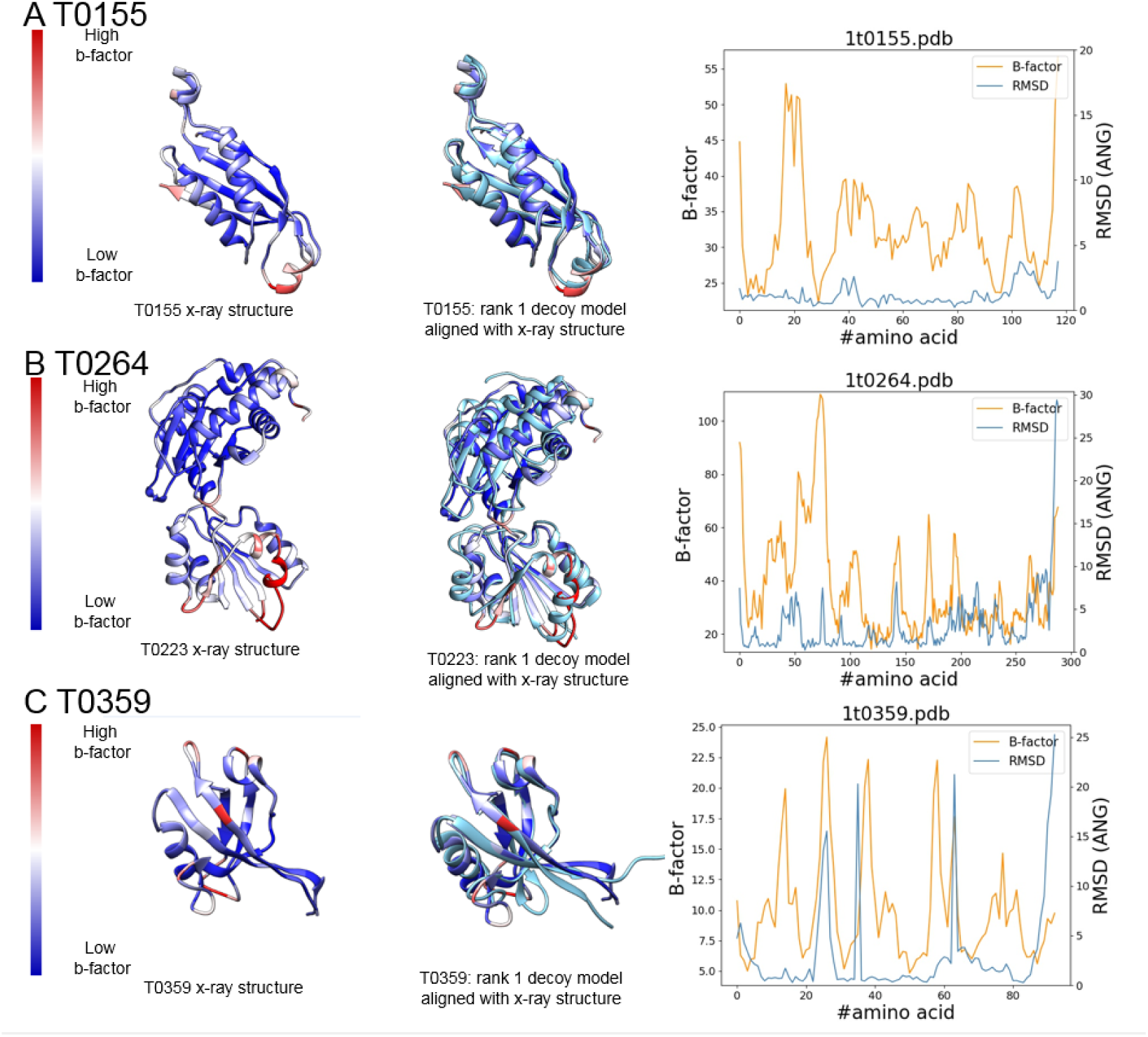
Relationship *between C*_***α***_ **RMSD values and the corresponding B-factors of the X-ray structures**. a) For T0155, the rank 1 structure picked by PBE have tm-score of 0.919 with its X-ray structure. b) For T0264, the rank 1 structure picked by PBE have tm-score of 0.847 with its X-ray structure. c) For T0359, the rank 1 structure picked by PBE have tm-score of 0.774 with its X-ray structure. The structure deviation mainly locates in the loop region or the terminal region.

These insights suggests that considering a prediction to be correct, only when there is an exact match between a predicted and an x-ray structure is not justifiable and unnecessary. For this reason, we decide to develop a new criterion to evaluate when a prediction can be considered to be correct.

Based on literature, we propose a new criterion where the TM-score between the predicted and the x-ray structures is used to evaluate the performance of the energy function. In detail, our novel evaluation criterion calculates the TM-score between the predicted structure with the lowest energy (rank one) and the x-ray structures. If this TM-score is above a certain threshold, the prediction is considered to be correct. Furthermore, the threshold is goal-oriented: If a virtually identical structure is required, the threshold could be 0.9, or if the prediction is used in drug design, the threshold could be 0.7, at which the average RMSD between predicted and x-ray structure is approximately 3Å (Figure 1).

## Performance on the novel criterion

We present the PBE performance on five data sets, when we use the novel TM-score criterion (TM-score > 0.7; table 2): On the CASP5-8, high resolution, and Rosetta (3D-Robot) datasets, the PBE achieves a success rate of 100%. Furthermore, on the Modeller_set (3d-robot) and I-TASSER_set (3d-robot) datasets, the success rates are 94.74% (only one of the predicted structures has a TM score of 0.66) and 97.44% (only one of the predicted structures has a TM score of 0.66), respectively. Overall, with the novel criterion, the PBE can be considered to be highly accurate.

**Table 2.**
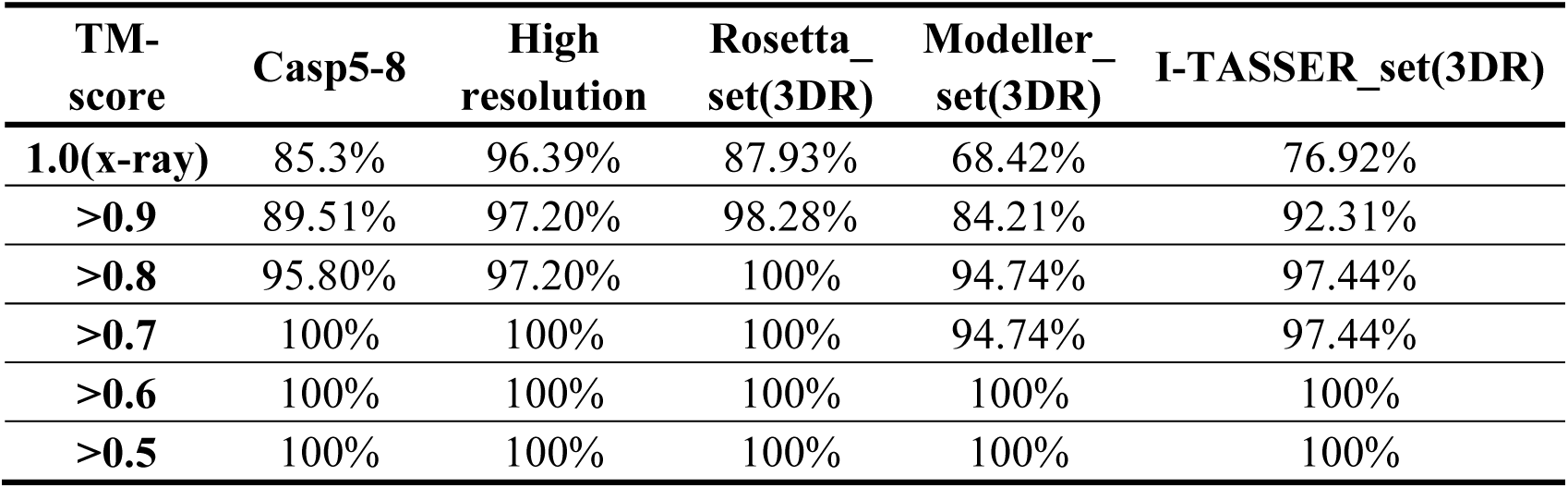
Percentage prediction success rate of the different TM-score threshold on independent datasets.

## Conclusion

To increase the accuracy of protein structure prediction, which is the key for its wider applications in the future, we present two significant improvements to the computational protein structure prediction. The first contribution is a novel knowledge-based Peptide-Bond-Interactions-based Energy (PBE) scoring function which predicts the lowest-energy structure from a set of possible (decoy) structures. The PBE scoring function not only outperforms the most-accurate, previously published statistical potential function, but also has much lower computational expense. In particular, the lower computational expense widens the PBEs potential usage. The second contribution is a novel criterion to evaluate the energy scoring function performance based on similarity with the x-ray structure instead of an exact match. This criterion is based on two realizations: (1) insignificant differences between predicted and x-ray structures lead to a prediction being classified as a failure, and (2) that the x-ray structure only represents an average native protein structure, due to the flexible nature of the protein structure. Using a combination of these two improvements, we achieved a 99.45% prediction success rate on all test datasets. Key to achieving the above results is to realize the importance of the peptide bond in the design of the protein energy function. Considering only the peptide bond is a good trade-off between prediction accuracy and computational expense.

## Method

### In-house dataset

The in-house dataset was obtained from the SCOP20 dataset^1^. The criteria for proteins included in the in-house dataset are: 1) its structure is not determined by NMR, and 2) the pair-wise sequence identity with those in independent test datasets is less than 20%. Finally, 5,147 proteins were contained in our in-house dataset. And for each protein, 400 random decoys were included using Rosetta de novo structure predictions followed by all-atom refinement^2^.

The template modeling score (TM-score)^3^ which measures the similarity of topologies of two protein structures was used as the criteria. TM-score has a range between (0, 1] with better similarity having higher TM-scores. And the values of the decoys ranged mainly between 0.2-0.4, as shown in the extended data figure 2.

### Independent datasets

Several of the widely used decoy datasets were employed as the independent ones to evaluate the performance of our new PBE, including:

### CASP5-8 dataset

The dataset was downloaded from https://zhanglab.ccmb.med.umich.edu/RW/^4^. All of the 143 proteins’ structures determined by X-ray and the corresponding decoy sets were remained for further use.

### High resolution dataset

The high resolution dataset including 148 proteins’ native structures and the decoy sets was downloaded from http://titan.princeton.edu/2010-10-11/HRdecoys/^5^. In the present work, only 107 protein structures determined by X-ray and the corresponding decoy sets were remained for further use.

### 3D-Robot dataset

The 3D-Robot decoy dataset was generated by the 3DRobot, a program devoted for automated generation of diverse and well-packed protein structure decoys. It includes structural decoys of the Rosetta, I-TASSER and Modeller sets, and was downloaded from https://zhanglab.ccmb.med.umich.edu/3DRobot/decoys/^6^. Among the proteins, 58, 39 and 19 X-ray determined structures and the corresponding decoy sets were remained for further use.

### Evolutionary information

Each protein’s evolutionary information was retrieved through applying the primary sequence to generate the position specific score matrix (PSSM) by running three iterations of PSI-BLAST against a non-redundant sequence database UniRef90 with an E-value cutoff of 0.001^7^. A 20-dimensinal vector *P* representing probabilities of conservation against mutations to 20 different amino acids was returned for each residue in the protein. And for a protein with sequence length *L*, the PSSM can be represented as an *L*\*20 matrix.

### Accuracy measures

We use two criteria to evaluate the performance accuracy: 1) after comparing with the X-ray structure, we present the number of structures regarded as rank 1 and that of those regarded within rank 5 that have the X-ray structure (this is the criterion typically used); 2) we list the number of rank 1 structures within the TM-score thresholds.

## Availability

The program named “pickingnative” has been written in C++ language and runs on the Linux platform. It is free of use on the web-server: www.166.111.152.74:8888/pbe_score/, and for academia user the executable code can be obtained after request by E-Mail.

## Author Contributions

Dr Xian-ming and Xia-yu have full access to all of the data in the study and take responsibility for the integrity of the data and the accuracy of the analysis.

*Study concept and design:* Xian-ming and Xia-yu

*Written all the computer programs*: Xian-ming

*Acquisition of data:* Xian-ming, Lu-yun, and Xia-yu

*Analysis and interpretation of data:* Xian-ming, Xia-yu, Lu-yun

*Drafting of the manuscript:* Xian-ming, Xia-yu, Lu-yun

*Critical revision of the manuscript for important intellectual content:* Xian-ming

*Statistical analysis:* Xian-ming, Xia-yu, Luyun

*Obtained funding:* Xian-ming.

*Administrative, technical, or material support:* Xian-ming.

*Study supervision:* Xian-ming

All authors have read the final manuscript.

## Disclaimer

The content is only the responsibility of the authors.

Correspondence authors Dr. Xian-ming Pan, or Dr. Xia-yu Xia School of Life Sciences, Tsinghua University, Beijing, 100084, China, Telephone: +86-10-62792827; E-Mail

## Extended data

**Extended Data Figure.1.**
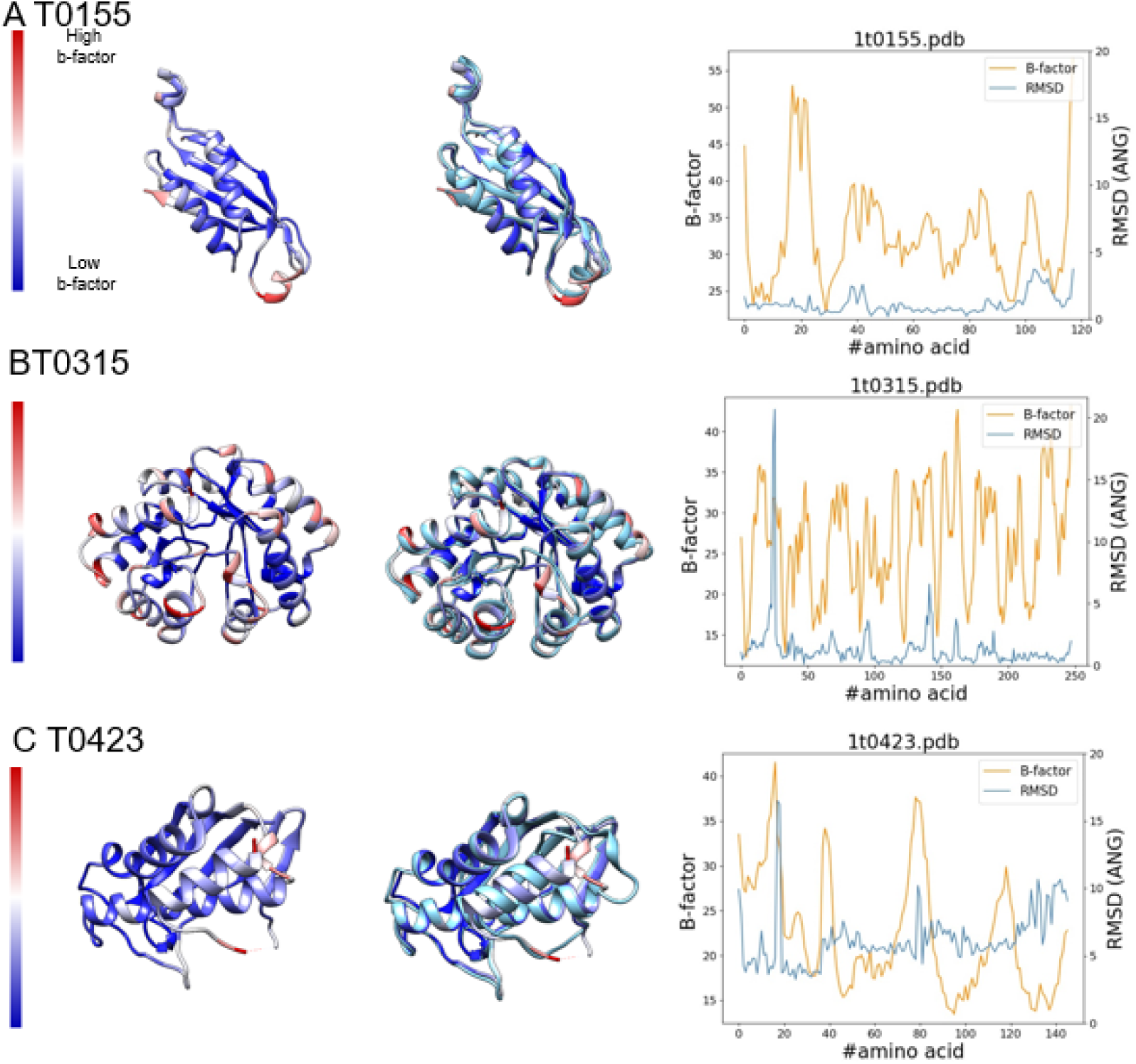
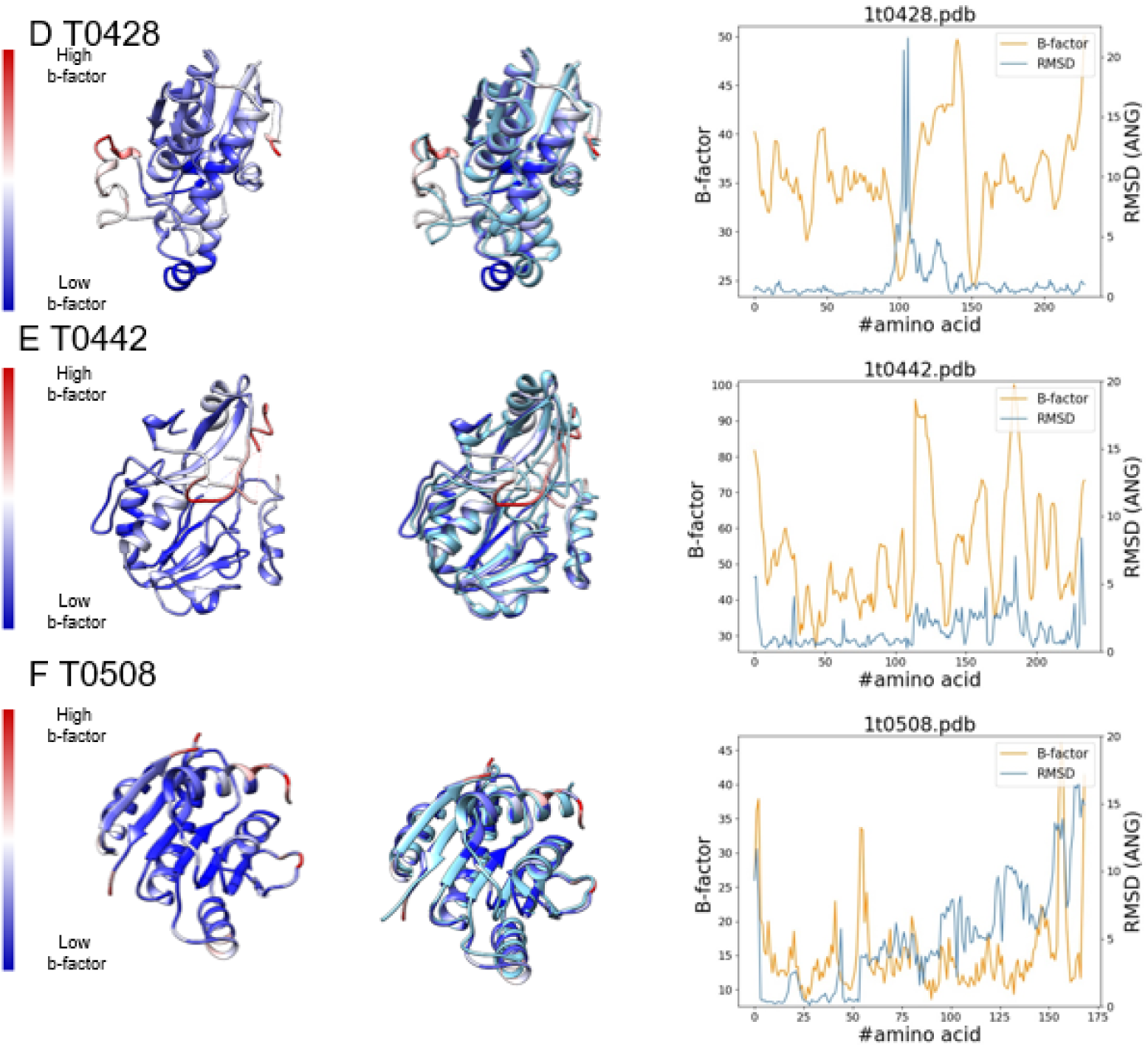
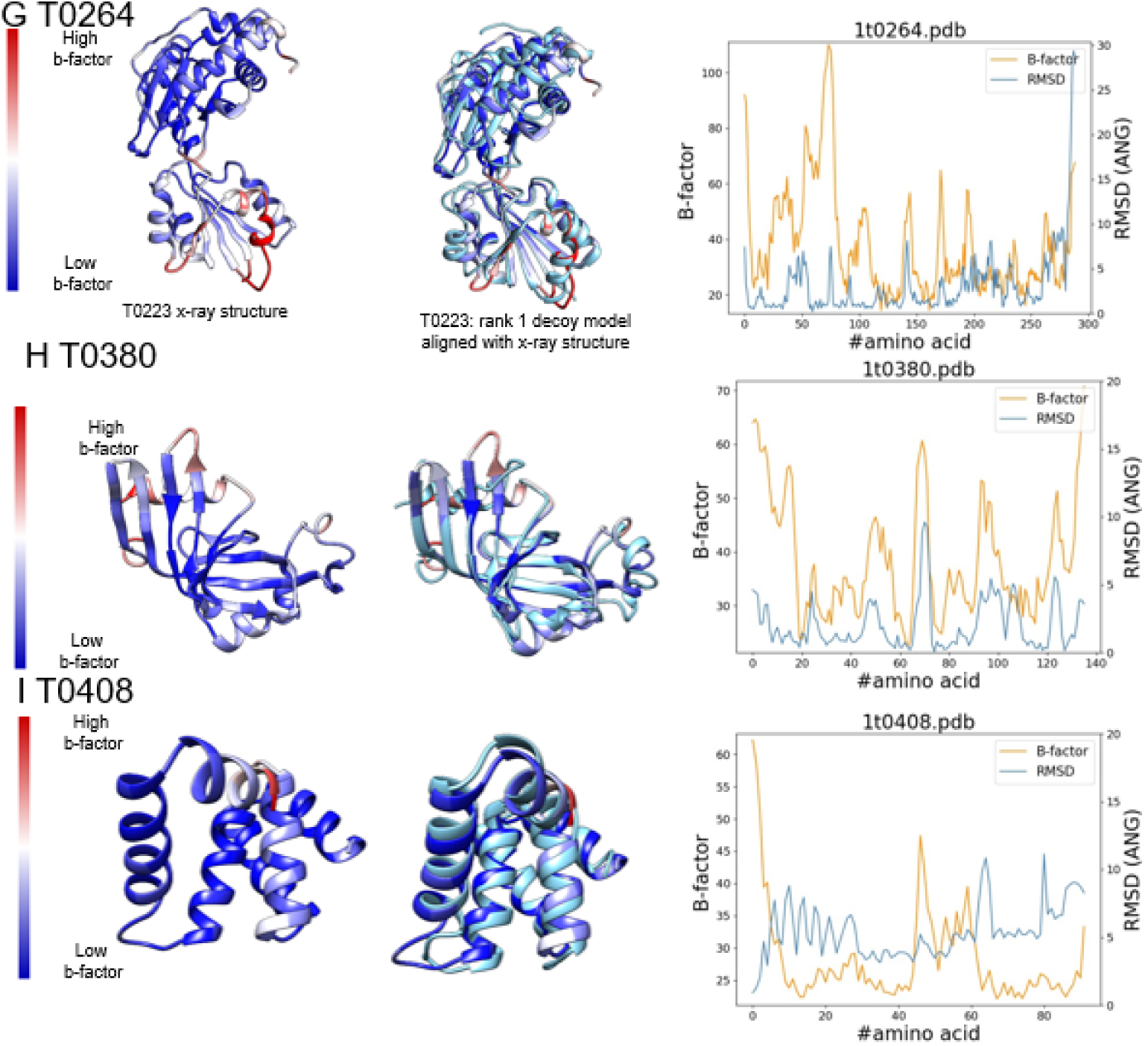
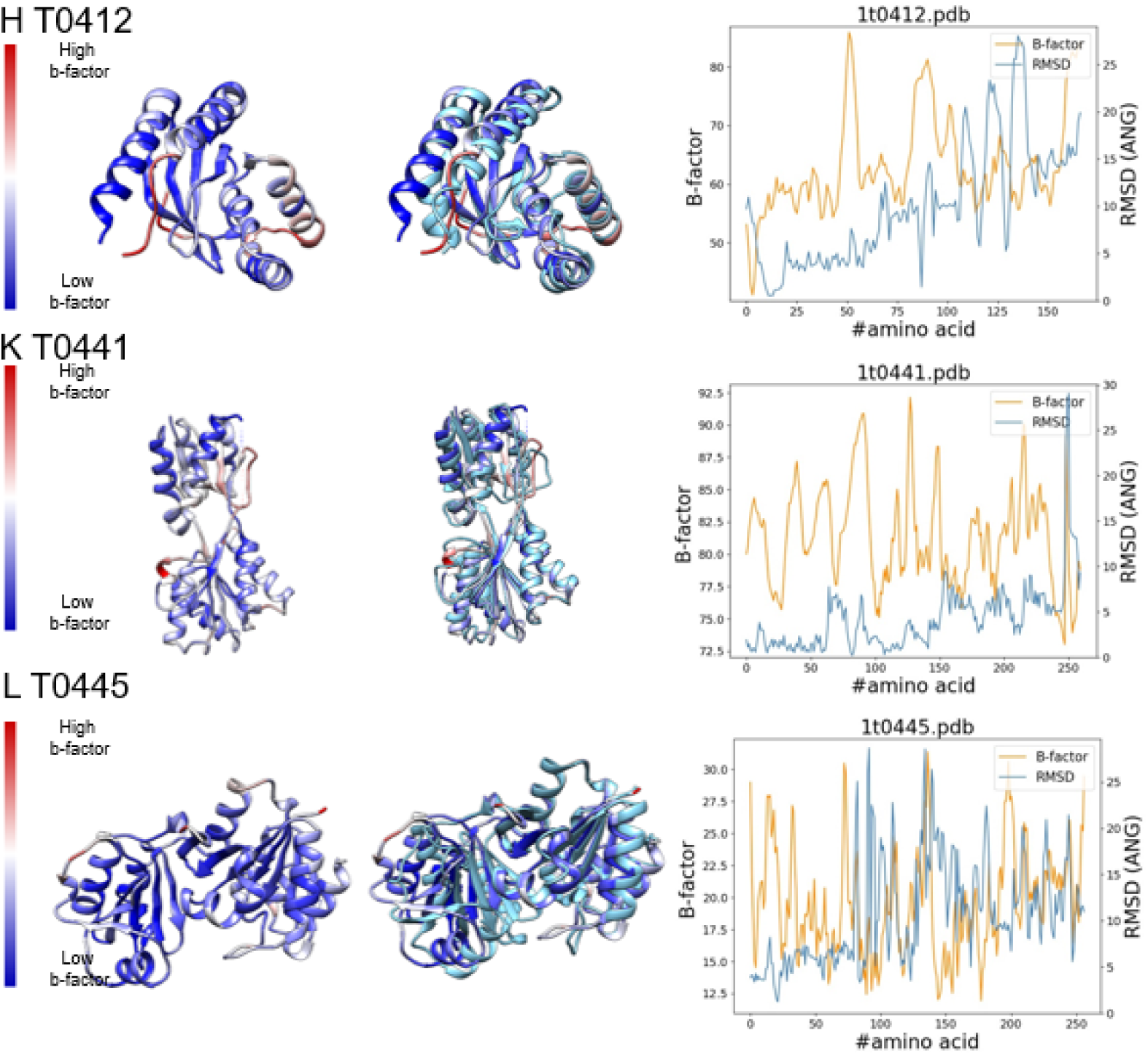
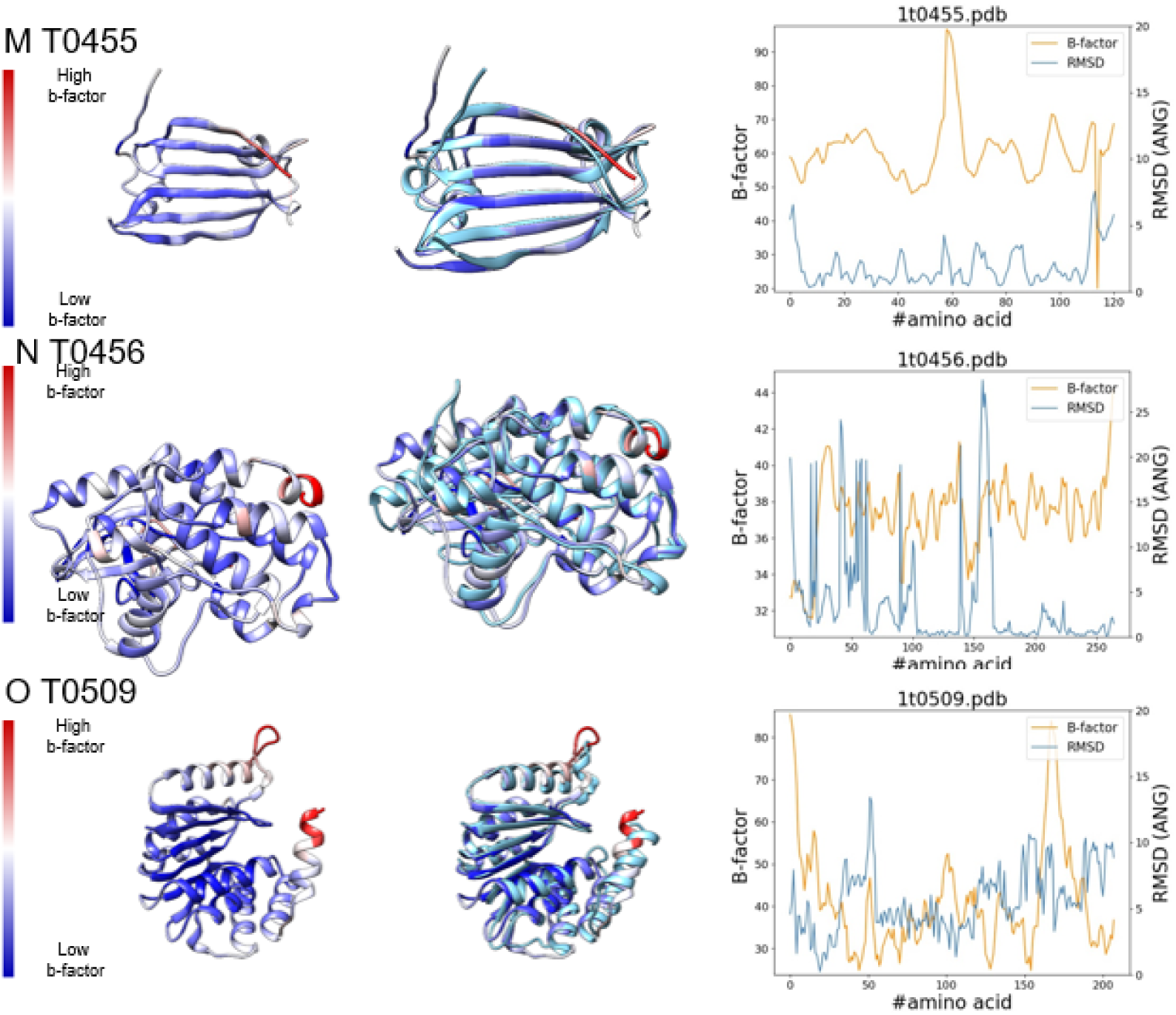
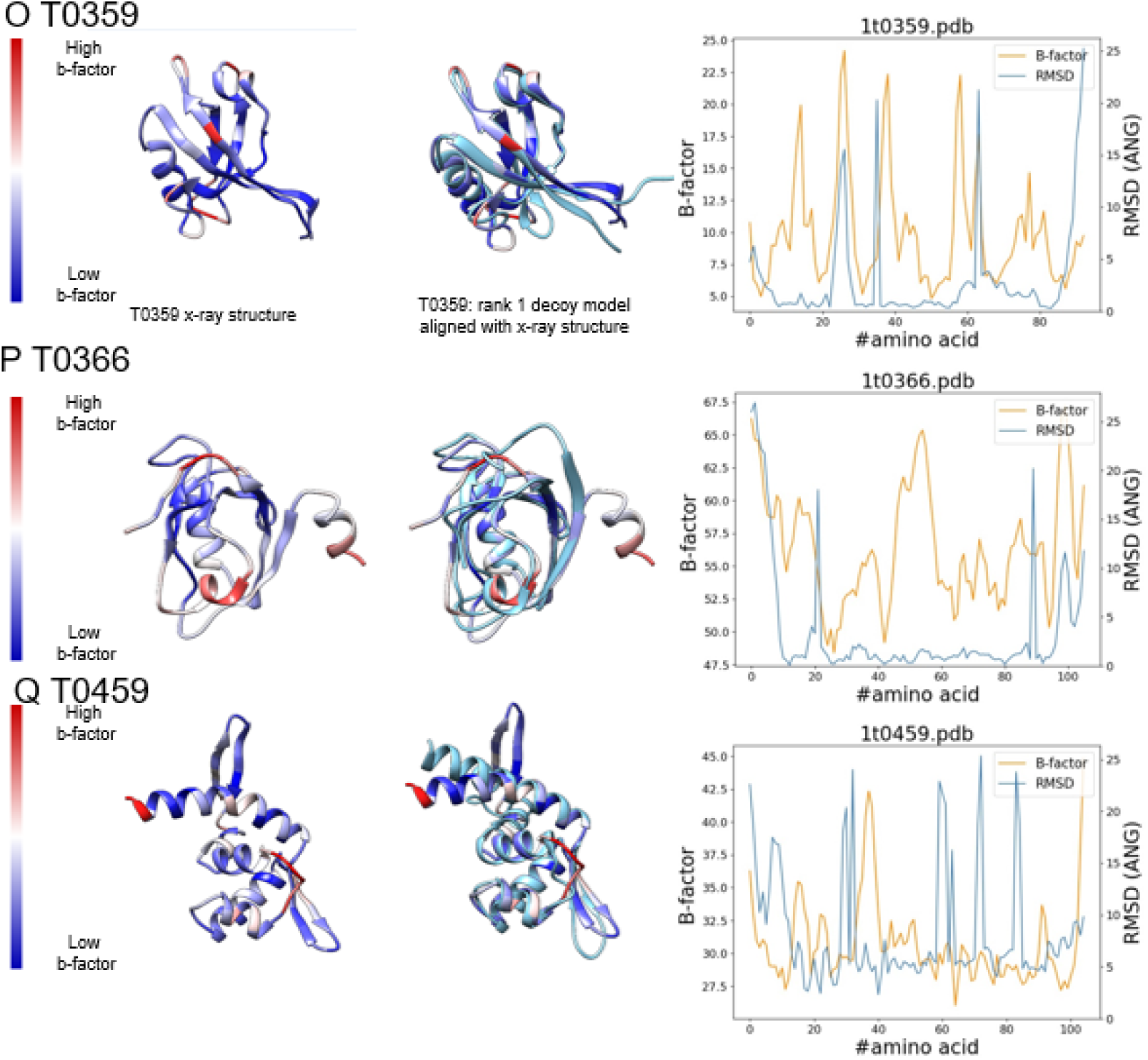
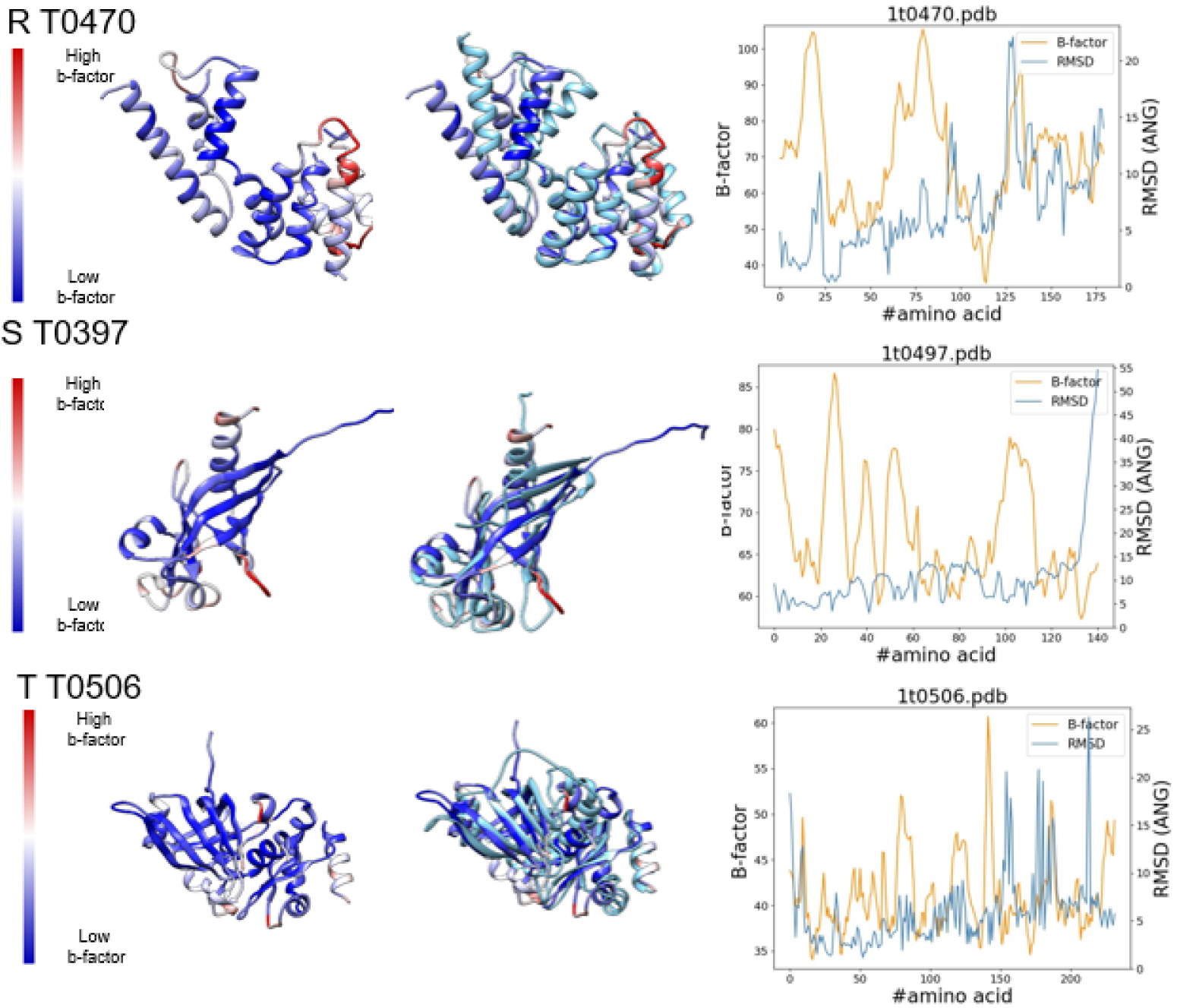
Relationship *between C*_***α***_ **RMSD values and the corresponding B-factors of the X-ray structures**.

**Extended Data Figure.2.**
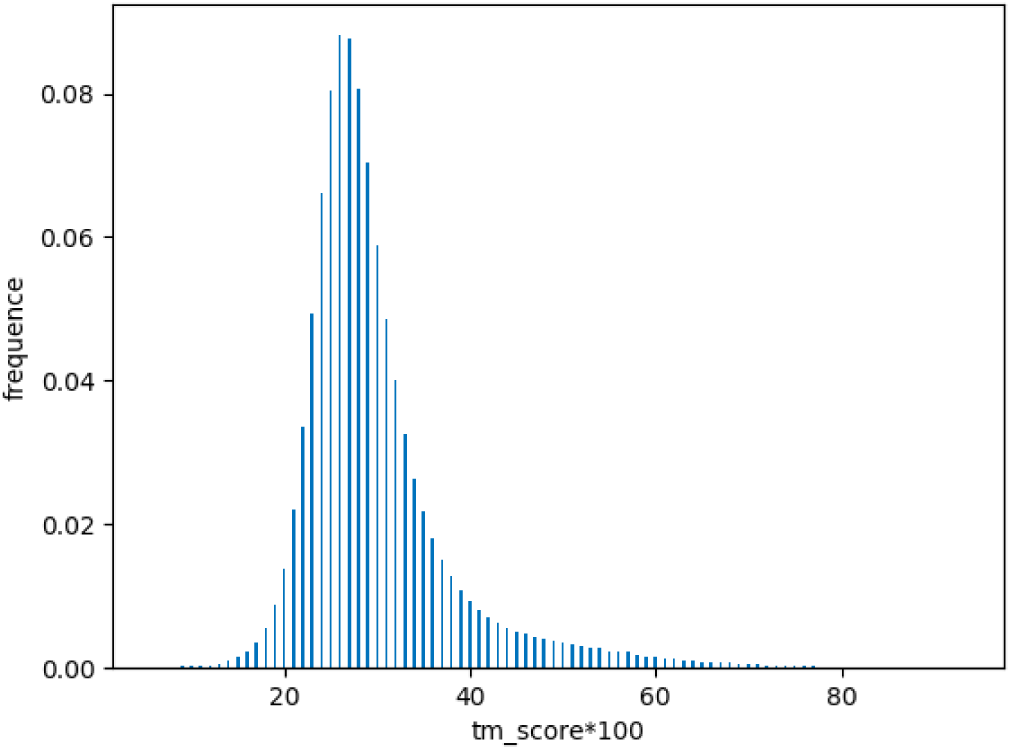
Distribution of the decoys’ *t*_***m***_ **scores of the in-house dataset**. The decoys’ TM-score mainly distribute in the range of 0.2-0.4

**Extended Data Table 1.** Selecting accuracy of the PBE score on inhouse dataset.

| TM-score | Self-consistency |  | Jackknife |  |
| --- | --- | --- | --- | --- |
|  | Accuracy / % | Z-score | Accuracy / % | Z-score |
| 1.0(x-ray) | 100.0 | -9.67 | 98.62 | -9.00 |
| >0.9 | 100.0 | -9.67 | 98.62 | -9.00 |
| >0.8 | 100.0 | -9.33 | 98.97 | -8.69 |
| >0.7 | 100.0 | -8.04 | 99.26 | -7.49 |
| >0.6 | 100.0 | -6.20 | 99.53 | -5.79 |
| >0.5 | 100.0 | -4.37 | 99.57 | -4.09 |

